# Early pandemic HIV-1 integration site preferences differ across anatomical reservoirs

**DOI:** 10.1101/2025.03.03.641190

**Authors:** Hinissan P. Kohio, Hannah O. Ajoge, Emile A. Barua, Neel R. Vajaria, Isaac K.F. Wu, Macon D. Coleman, Sean K. Tom, Frank van der Meer, John Gill, Deirdre Church, Paul Beck, Christopher Power, Guido van Marle, Stephen D. Barr

## Abstract

Despite effective viral suppression with modern antiretroviral therapy (ART), HIV-1 persists in latent reservoirs across multiple tissues. Integration into the host genome is essential for viral persistence, yet the characteristics of these reservoir sites across anatomical locations remain poorly understood. To address this, we analyzed integration sites from matched esophagus, PBL/PBMC, stomach, duodenum, colon, and unmatched brain tissue samples of individuals infected with HIV-1 subtype B. The virus used in this study was from 1993, an early stage of the HIV pandemic, providing insights into integration patterns before extensive ART use. Our analysis examined genomic feature enrichment, proximity to non-B DNA structures, integration hotspots, and site overlap across tissues and individuals. We identified a distinct integration pattern in brain tissue, characterized by reduced gene targeting and increased enrichment in Short Interspersed Nuclear Elements (SINEs) and DNase I hypersensitivity sites (DHS). Tissue-specific preferences for integration near non-B DNA structures were evident, alongside shared and unique hotspots across tissues and individuals. Notably, genes associated with HIV-related diseases were frequent integration targets. These findings underscore the complex interplay between viral integration, host genetics, and tissue-specific factors, highlighting the potential role of integration sites in disease development.

## Introduction

The widespread use of antiretroviral therapy (ART) has significantly improved the quality of life for individuals living with HIV-1. While ART effectively suppresses plasma viremia to undetectable levels, it is not curative due to the persistence of viral reservoirs—latently infected cells that can reactivate upon treatment interruption (1–3). These reservoirs consist of cell types or anatomical sites where replication-competent virus persists longer than the actively replicating viral population (4–6). During early infection, HIV-1 preferentially integrates into transcriptionally active regions of the genome to maximize proviral expression and viral dissemination. However, this strategy comes with risks: the production of viral proteins can trigger immune-mediated cell clearance or virus-induced cytopathic effects. To evade immune detection and promote long-term persistence, HIV-1 often integrates into transcriptionally inactive regions of the host genome, limiting viral gene expression and facilitating immune evasion. These transcriptionally repressive genomic regions are characterized by distinct features. For instance, lamina-associated domains (LADs) create a chromatin environment that is tightly linked to the nuclear periphery, suppressing transcription (7, 8). Short interspersed nuclear elements (SINEs) and other transposable elements contribute to gene silencing through repressive chromatin marks (9, 10). Additionally, non-B DNA structures—such as G-quadruplexes (G4), cruciform DNA, triplex DNA, and Z-DNA—can modulate transcription by interfering with gene regulatory mechanisms (11–21). Recent studies have shown that in both elite controllers and individuals on long-term ART, most integrated HIV-1 proviruses are located in these transcriptionally repressive genomic features, reinforcing the role of integration site selection in viral persistence (8, 22–30).

Despite extensive research on blood-based HIV-1 reservoirs, integration site profiling across diverse anatomical tissues remains limited. HIV-1-infected cells persist in a wide range of tissues, including the brain, lungs, kidneys, liver, adipose tissue, lymphoid organs, gastrointestinal tract, male and female genitourinary systems, and bone marrow (31). While CD4+ T cells serve as the primary latent reservoir (1, 32–35), macrophages and organ-specific cells—such as epithelial cells, microglia, astrocytes, and podocytes—also harbor integrated virus (36–38). Myeloid cells, in particular, have been increasingly recognized as significant reservoirs (39). A recent study by Wu et al. (2020) assessed HIV-1 integration sites in sorted memory, tissue-resident, and follicular helper CD4+ T cell subsets from paired peripheral blood mononuclear cells (PBMCs) and lymphoid tissues (inguinal, cervical, and tonsil) (40). They found that HIV-1 reservoirs exhibited tissue compartmentalization, but integration site patterns were largely conserved across anatomical sites, cell subsets, and infection stages, with minimal overlap between different compartments.

To further elucidate the heterogeneity of tissue-specific HIV-1 reservoirs, we characterized the genomic environment of integrated proviruses from a unique biobank of early-pandemic HIV-1 subtype B-infected individuals. Our analysis included matched esophagus, PBL/PBMC, stomach, duodenum, and colon tissues, along with unmatched brain tissue. We identified distinct integration site profiles across these anatomical sites, with substantial overlap of integration sites and hotspots across tissues and individuals. These findings provide new insights into how integration site selection influences HIV-1 persistence and may inform future strategies to target and disrupt viral reservoirs.

## RESULTS

### Brain tissue exhibits a distinct HIV-1 integration profile with reduced gene targeting and increased SINE enrichment

Most HIV-1 integration studies have focused on PBMCs or blood-derived CD4+ T cells, with limited characterization of tissue-specific integration patterns. Given the compartmentalization of HIV-1 *in vivo* and the potential influence of tissue residency on integration site selection, we sought to address this knowledge gap (40, 41). We analyzed integration site profiles from patient-matched tissue samples, including esophagus (165 integration sites), PBL/PBMC (161 sites), stomach (202 sites), duodenum (137 sites), and colon (119 sites) from five individuals infected with HIV-1 subtype B between 1993 and 2010, all of whom were receiving antiretroviral therapy (Table S1). Additionally, we examined 145 integration sites from brain tissue samples obtained from a separate, unmatched cohort of eight HIV-1 subtype B-infected individuals who were not receiving ART and had succumbed to AIDS-related illnesses. Integration site mapping was performed using an established in-house pipeline (28–30).

Across all tissues, we identified 929 unique integration sites, excluding those resulting from clonal expansion or mapping to ambiguous genomic regions. To assess tissue-specific preferences, we compared the distribution of integration sites across various genomic features—including CpG islands, DNase I hypersensitivity sites (DHS), endogenous retroviruses (ERVs), heterochromatin, SINEs, LINEs, low-complexity regions (LCRs), oncogenes, genes, simple repeats, and transcription start sites (TSS)— against a matched random control (MRC). Figure 1A summarizes these integration patterns, highlighting largely consistent site preferences across tissues, with a few notable exceptions (Table S2). For instance, the duodenum exhibited enrichment in CpG islands and LCRs, while both the duodenum and brain showed preferential integration within oncogenes. Notably, brain tissue displayed significant enrichment in SINEs and DHS.

**Figure 1:**
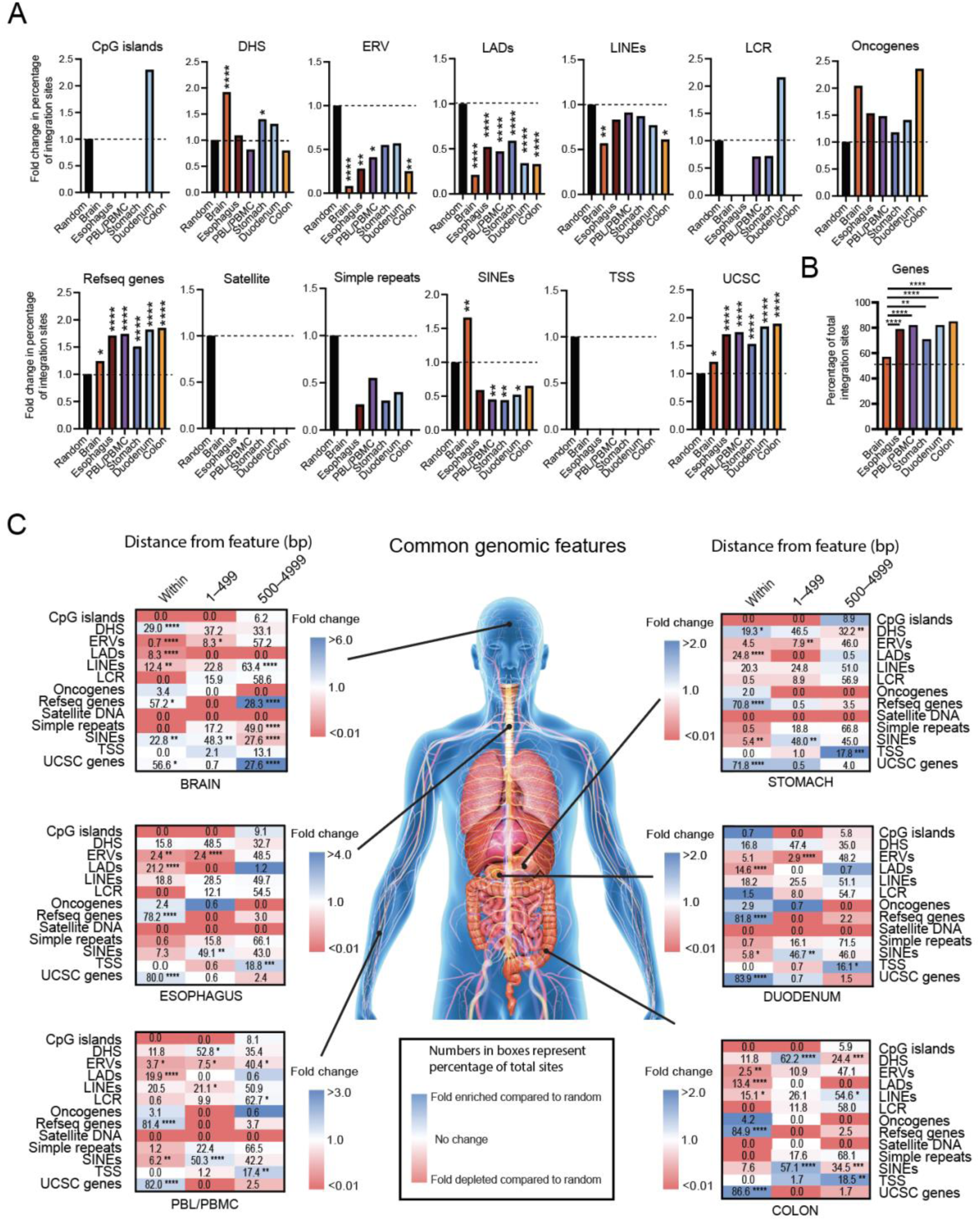
Integration site preference for common genomic features differ in brain compared to other tissues. A, Comparisons of the fold-change in the percentage of integration sites located within various common genomic features from brain (n= 145 sites), esophagus (n= 165 sites), PBL/PBMC (n= 161 sites), stomach (n= 202 sites), duodenum (n= 137 sites), and colon (n= 119 sites) (total sites= 929). Statistical comparisons are made with respect to the matched random control (MRC). B, Comparison of the fold-change in the percentage of integration sites located within genes of the different tissues. Statistical comparisons are made with respect to brain. C, Heatmaps illustrating the fold-enrichment or depletion of unique integration sites (compared to the MRC) within, 1-499 bp or 500-4,999 bp away from the feature. Darker shades represent higher fold-changes in the ratio of integration sites to MRC sites. Numbers inside the boxes represent the percentage of total sites located in that bin. Significant differences are denoted by asterisks; *P < 0.05, **P < 0.01, ***P < 0.001, ****P < 0.0001) (Chi-square test with Yates’ correction).

A key distinction emerged in gene targeting patterns. While integration within genes was consistently high across PBL/PBMC (82%), esophagus (79%), stomach (71%), duodenum (82%), and colon (85%), brain tissue exhibited significantly lower gene targeting, with only 57% of integration sites occurring within genes (Figure 1B, Table S2). For comparison, 46% of integration sites in the MRC dataset were within genes, suggesting that brain integration deviates substantially from other tissues. To further dissect spatial relationships between integration sites and genomic features, we classified sites into three distance bins: within the feature, 1–499 bp away, and 500–4,999 bp away (Figure 1C, Table S2). Compared to the MRC, integration preferences were largely conserved across most tissues, except for brain. Brain-derived integration sites were uniquely enriched 500–5,000 bp away from genes, whereas all other tissues showed enrichment within 1–499 bp of DHS and 500–5,000 bp of TSS.

Together, these findings indicate that while most tissues share common integration site preferences, brain tissue exhibits a distinct profile characterized by reduced gene targeting and preferential integration into SINEs. This divergence suggests that brain-specific chromatin environments may play a role in shaping HIV-1 persistence within the central nervous system.

### Distinct HIV-1 integration patterns near non-B DNA across anatomical tissues

To investigate tissue-specific preferences for HIV-1 integration near non-B DNA structures, we analyzed integration sites within 500 bp of various non-canonical DNA motifs, including A-phased DNA, cruciform structures, G-quadruplexes (G4), inverted repeats, triplex DNA, Z-DNA, and others. This 500 bp window was selected based on its potential functional impact on transcriptional regulation (29). As shown in Figure 2A, integration preferences varied by tissue. Brain-derived sites were less frequently associated with A-phased DNA, cruciform structures, G4, inverted repeats, triplex DNA, and Z-DNA but showed modest enrichment near direct repeats and slipped DNA. Esophageal sites followed a similar pattern, with reduced association near A-phased DNA, cruciform structures, slipped DNA, triplex DNA, and Z-DNA, while showing modest enrichment near G4 and short tandem repeats (STRs). PBL/PBMC sites exhibited modest enrichment near G4, mirror repeats, and STRs. Stomach-derived sites favored mirror repeats and Z-DNA, whereas duodenum sites were enriched near cruciform structures and mirror repeats but disfavored A-phased DNA, direct repeats, slipped DNA, triplex DNA, and Z-DNA. Colon sites showed enrichment near G4, mirror repeats, and STRs, while exhibiting lower association with cruciform structures, triplex DNA, and Z-DNA.

**Figure 2:**
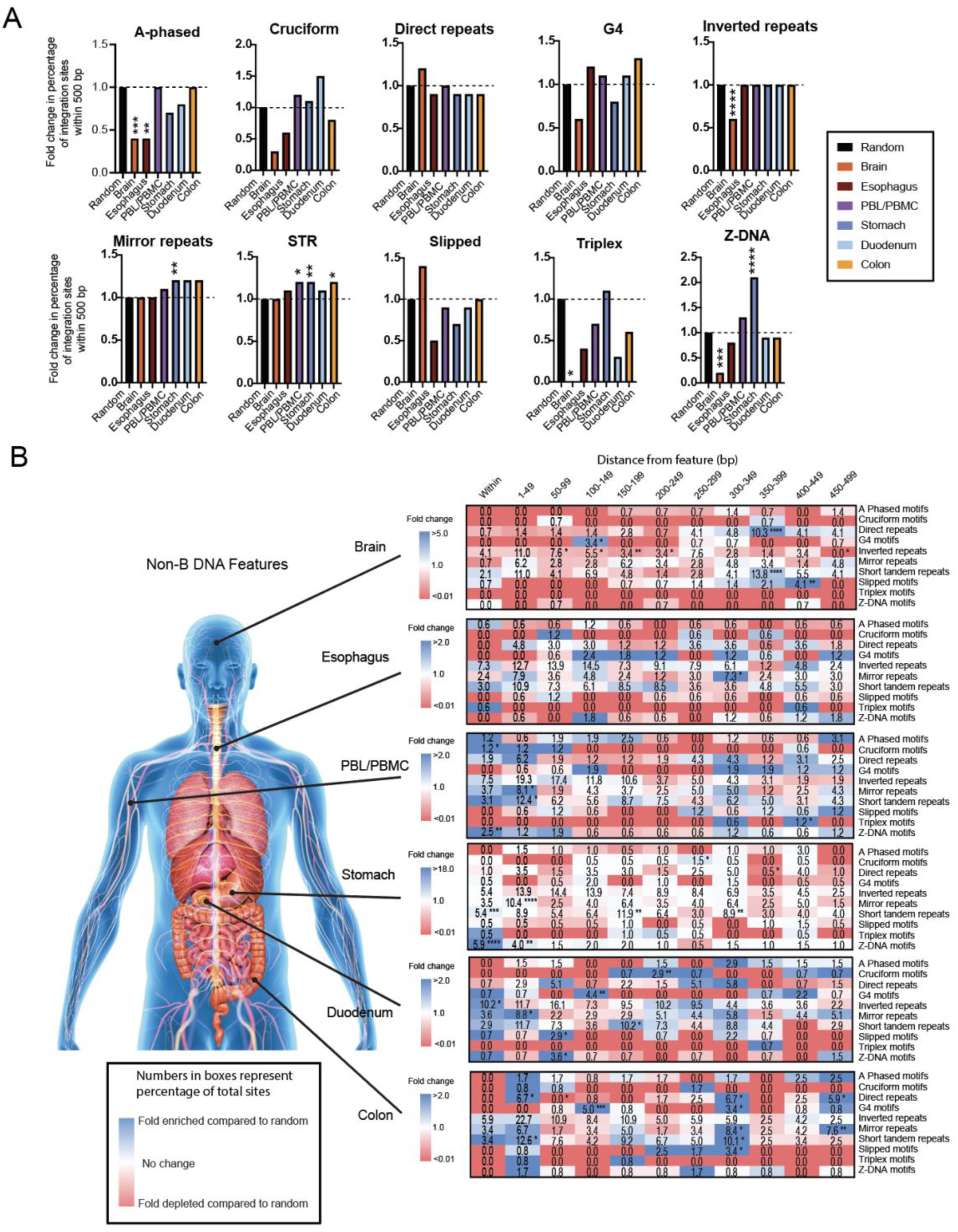
Integration site preference for non-B DNA features differ across tissues. A, Comparisons of the fold-change in the percentage of integration sites located within 500 bp of various non-B DNA features from brain (n= 145 sites), esophagus (n= 165 sites), PBL/PBMC (n= 161 sites), stomach (n= 202 sites), duodenum (n= 137 sites), and colon (n= 119 sites) (total sites= 929). Statistical comparisons are made with respect to the matched random control (MRC). B, Heatmaps illustrating the fold-enrichment or depletion of unique integration sites compared to the MRC. Each bin represents sites located: within, 1-49 bp, 50-99 bp, 100-149 bp, 150-199 bp, 200-249 bp, 250-299 bp, 300-349 bp, 350-399 bp, 400-449 bp, or 450-499 bp away from the feature. Darker shades represent higher fold-changes in the ratio of integration sites to MRC sites. Numbers inside the boxes represent the percentage of total sites located in that bin. Significant differences are denoted by asterisks; *P < 0.05, **P < 0.01, ***P < 0.001, ****P < 0.0001) (Chi-square test with Yates’ correction).

Given our prior findings that HIV-1 integration preferences near non-B DNA vary with distance (28–30), we next examined integration patterns using 50 bp sliding windows (Figure 2B, Table S3). This analysis revealed distinct distance-dependent tissue-specific preferences: Brain: HIV-1 integration was enriched at greater distances from several non-B DNA features, including G4 (100–149 bp, 4.4-fold, 3.4% of sites, P < 0.05), direct repeats (350–399 bp, 4.6-fold, 10.3% of sites, P < 0.0001), STRs (350– 399 bp, 3.0-fold, 13.8% of sites, P < 0.0001), and slipped motifs (400–449 bp, 5.8-fold, 4.1% of sites, P < 0.01). PBL/PBMC: These sites showed a stronger tendency for integration within or near most non-B DNA features, particularly Z-DNA (14.2-fold, 2.5% of sites, P < 0.01), mirror repeats (2.0-fold, 8.1% of sites, P < 0.05), and cruciform structures (9.5-fold, 1.2% of sites, P < 0.05). Stomach: HIV-1 integration was highly enriched within Z-DNA (18-fold, 5.9% of sites, P < 0.0001) and STRs (3.7-fold, 5.4% of total sites, P < 0.001), with additional enrichment near mirror repeats (1–49 bp, 2.7-fold, 10.4% of total sites, P < 0.0001). Duodenum: This tissue showed significant enrichment near inverted repeats (1.9-fold, 10.2% of total sites, P < 0.05) and mirror repeats (2.0-fold, 8.8% of total sites, P < 0.05), while HIV-1 integration sites were positioned further from G4, Z-DNA, cruciform structures, and slipped motifs. Colon: Integration sites were enriched near direct repeats (2.7-fold, 6.7% of total sites, P < 0.05) and STRs (1.9-fold, 12.6% of total sites, P < 0.05). Additionally, HIV-1 integration sites tended to be located further from G4, direct repeats, mirror repeats, STRs, and slipped motifs (300–349 bp, 2.3–4.5-fold, 3–10% of sites, P < 0.05). These findings highlight the tissue-specific nature of HIV-1 integration site selection relative to non-B DNA structures, suggesting that chromatin organization and DNA topology may influence viral persistence in different anatomical compartments.

### Integration sites overlap across tissues and individuals

We examined the extent to which HIV-1 integration sites were shared among different tissues. Out of 929 unique integration sites, 159 (17.1%) were found in more than one tissue (Figure 3A, Table S4). Most of these overlaps involved two tissues (70 sites, 7.5%), followed by three tissues (53 sites, 5.7%), and four tissues (26 sites, 2.8%). A smaller number of sites overlapped across five (6 sites, 0.6%) and six (4 sites, 0.4%) tissues. Among the most frequent overlaps: 13 sites were shared between PBL/PBMC, duodenum, and stomach; 11 sites were found in PBL/PBMC, duodenum, stomach, and colon; 11 sites were shared between the esophagus and stomach; and 11 sites were found in the esophagus, stomach, and duodenum. This observed overlap contrasts sharply with the matched random dataset, where only one overlapping site (between PBL/PBMC and stomach) was detected out of 13,055 total sites (<0.0001%). This reconfirms that HIV-1 integration is non-random and that certain genomic regions may be preferentially targeted across multiple tissues.

**Figure 3:**
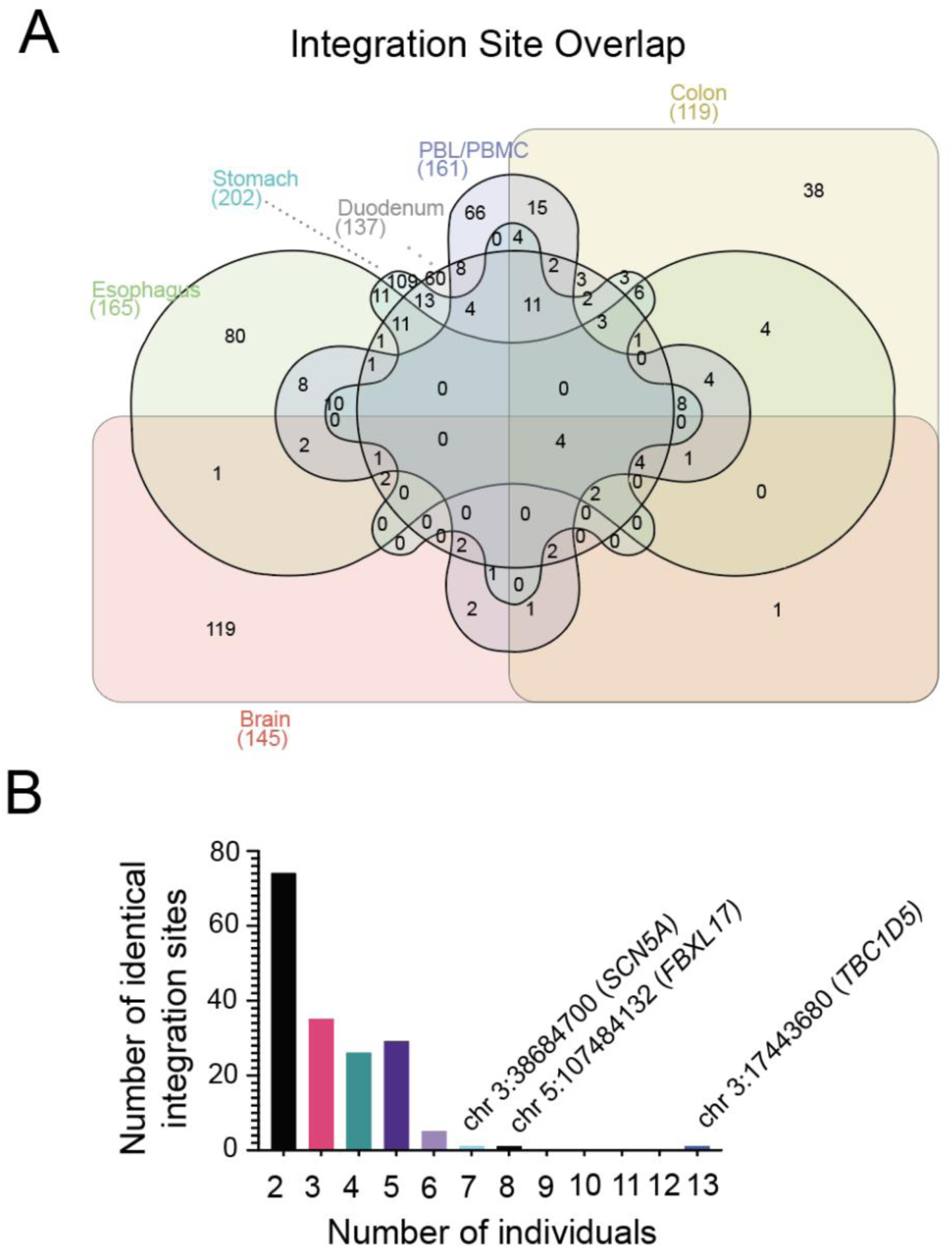
Integration site overlap. A, Venn diagram showing the number of identical integration sites shared across tissues. B, Comparison of the number of identical integration sites between a number of different individuals. Inset, chromosomal location of the single overlapped integration site in those individuals.

To determine whether integration site overlap extended across individuals, we analyzed shared sites among the five individuals with matching PBL/PBMC, esophagus, stomach, duodenum, and colon samples, as well as the eight individuals with brain tissue (13 individuals total). Of the 929 unique integration sites, 172 (18.5%) were found in more than one individual (Figure 3B, Table S4). Most overlapping sites were shared between two individuals (74 sites), with smaller numbers found across three (35 sites), four (26 sites), five (29 sites), six (5 sites), seven (1 site), and eight (1 site) individuals. Notably, one integration site (chr3:17443680, within the *TBC1D5* gene) was present in all 13 individuals. Additional recurrent sites included *FBXL17* (chr5:107484132), found in eight individuals, and *SCN5A* (chr3:38684700), found in seven individuals. These findings highlight significant HIV-1 integration site overlap, both across different tissues within individuals and across individuals, suggesting that certain genomic loci may serve as preferential sites for viral integration.

### Integration site hotspots exist across tissues

To identify regions of frequent HIV-1 integration, we defined hotspots as genomic loci where multiple integration events occurred within 100 bp of each other. Brain tissue exhibited the highest proportion of integration hotspots, with 45% of total integration sites clustering within these regions—significantly more than all other tissues (P < 0.0001). Other tissues also displayed notable hotspot integration, including esophagus (21%), duodenum (15%), colon (10%), stomach (10%), and PBL/PBMC (8%) (Figure 4A, Table S4). Some hotspot integrations were observed at identical chromosomal positions across different tissues (Figure S1, Table S4), suggesting recurrent targeting of specific genomic loci. While the chromosomal distribution of integration sites was largely similar among tissues, a striking exception was observed in brain tissue: chromosome 9 accounted for 28% of all brain-derived integration sites, compared to only 5–10% in other tissues (Figure 4B-4D, Table S4). This was driven by a dense cluster of 39 brain-derived integration sites within a 313 bp window on chromosome 9 (chr9:129,604,522–129,604,835) (Figure 4C).

**Figure 4:**
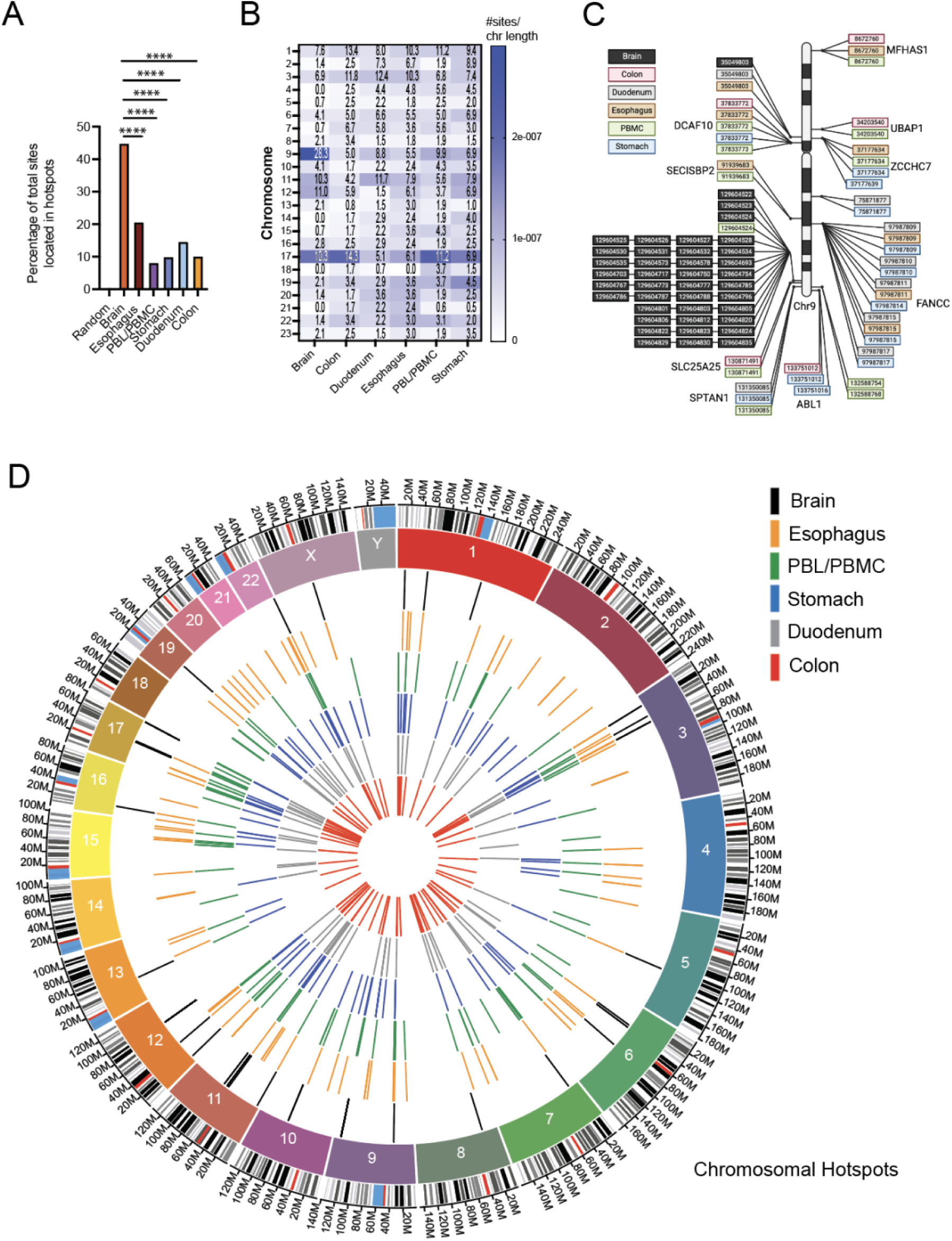
Chromosomal hotspots for integration. A, Comparison of the percentage of total integration sites located within 100 bp of each other (‘hotspots’). Statistical comparisons are made with respect to brain. Significant differences are denoted by asterisks; *P < 0.05, **P < 0.01, ***P < 0.001, ****P < 0.0001) (Chi-square test with Yates’ correction). B, Comparison of the percentage of total integration sites (numbers inside boxes on the heatmap) on each chromosome within each tissue. Blue shading represents the normalized number of integration sites per chromosome length. C, Schematic showing the approximate locations of integration site hotspots on chromosome 9 from brain. If integration hotspots are located within a gene, the gene name is provided adjacent to the integration site cluster. D, Circos plot showing the approximate locations of integration site hotspots on each chromosome from each tissue.

We next examined which genes were most frequently targeted for integration across tissues. The total number of genes harboring integration sites varied by tissue: brain, 70 genes; PBL/PBMC, 129 genes; esophagus, 107 genes; stomach, 134 genes; duodenum, 99 genes; and colon, 97 genes (Figure 5A, Table S5)(42). Many integration sites were tissue-specific, occurring in genes uniquely targeted within a given tissue: brain, 42 genes; PBL/PBMC, 38 genes; esophagus, 35 genes; stomach, 54 genes; duodenum, 33 genes; and colon, 20 genes (Figure 5B, Table S5). Despite this tissue specificity, certain genes were recurrent integration hotspots, with five or more unique integration events: brain, *ATP2C1*, *CHKA*, *ST3GAL3*; PBL/PBMC, *SYNRG*; esophagus, *MBOAT7*; stomach, *ANKFY1*, *CARD8*; duodenum, *DEPDC5* (Figure 5B, Table S5). Notably, some genes were integration hotspots across multiple tissues. Four tissues shared *ASCC1*, *SSH2*, *ZCCHC11*, *ZCCHC7*, *CCNL2*, *MAP3K3*, *MFSD11*, and *PTPRD*. Five tissues shared *TBC1D5*, *EIF4G3*, *PPP6R2*, and *RBM6*. The gene *SCN5A* was a recurrent integration hotspot in all six tissues (Figure 5C, Table S5).

**Figure 5:**
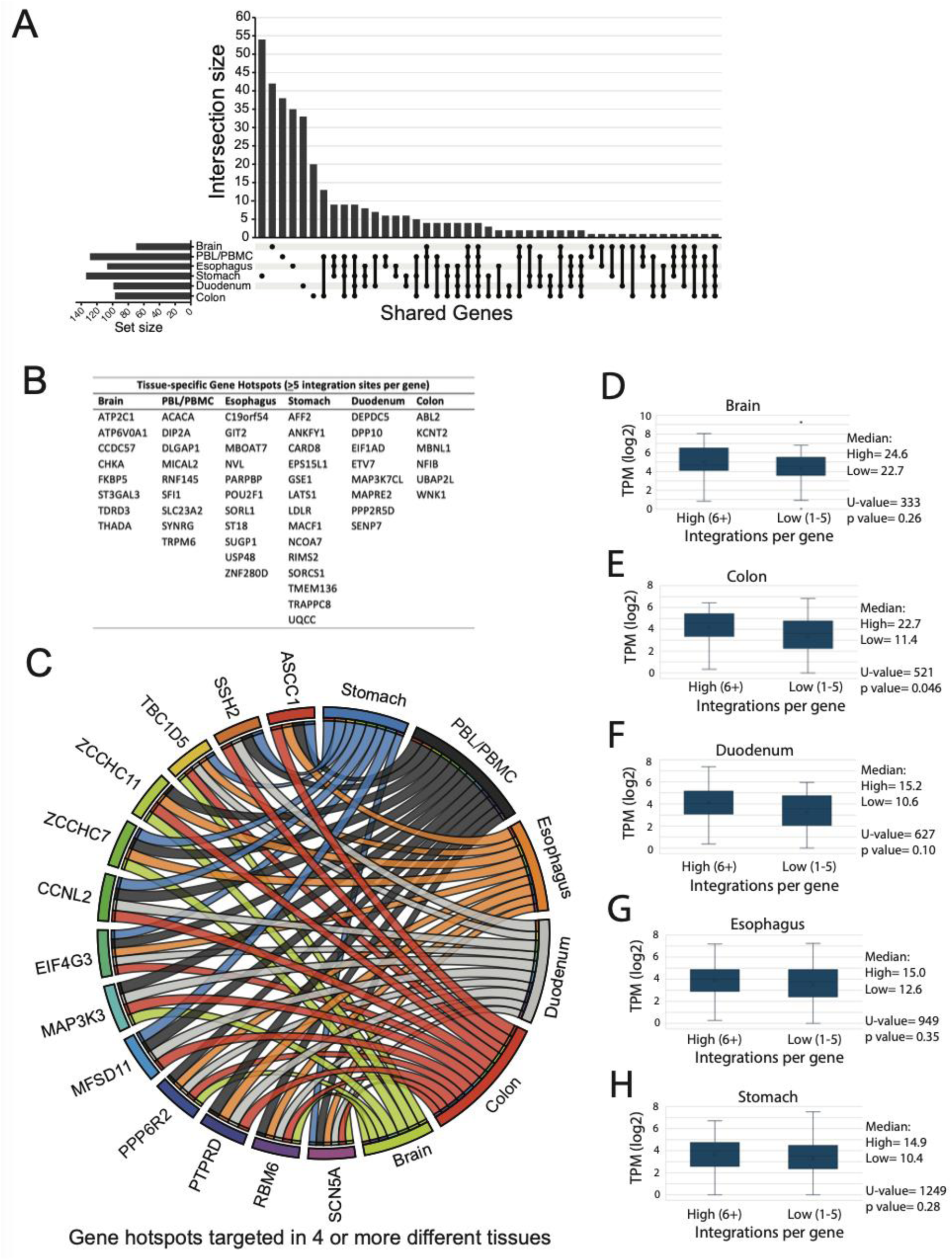
Integration site gene hotspots across tissues. A, UpSet plot displaying the intersection of gene sets hosting HIV-1 integration sites located within 100 bp of each other (“gene hotspots”) across different tissues. The horizontal bars represent the total number of gene hotspots identified in each tissue, while the vertical bars indicate the number of shared gene hotspots among tissue-specific sets. Black dots and connecting lines denote specific intersections, with single dots representing unique hotspots within one tissue and multiple connected dots indicating shared hotspots across tissues. The dataset was filtered to retain only genes with integration sites meeting the 100 bp proximity threshold, and intersection sizes reflect the degree of conservation or tissue specificity of integration patterns. B, All hotspot genes hosting ten or more integration sites (‘gene super-hotspots’) were filtered for each tissue and compared to each other to identify genes highly targeted by four or more tissues. The ribbons emerging from each tissue in the Circos plot connect to the genes shared by the tissues. C, Tissue-specific gene hotspots hosting five or more integration sites per gene are listed. These represent genes not targeted by any of the other tissues. D-H, Box-and-whisker plots display the distribution of gene expression levels (transcripts per million, TPM) for two groups of genes: those hosting six or more HIV-1 integration sites (high-frequency group) and those hosting 1 to 5 integration sites (low-frequency group). Gene expression data were obtained from RNA-seq datasets available in the Human Protein Atlas for the brain, colon, duodenum, esophagus, and stomach. The central line within each box represents the median expression level, the box spans the interquartile range (IQR), and whiskers extend to 1.5 times the IQR. Outliers are represented as individual points. Statistical significance was assessed using a Mann-Whitney U Test (Wilcoxon Rank-Sum Test) to determine whether gene expression levels significantly differ between the two integration frequency groups across tissues.

To assess whether HIV-1 preferentially integrates into highly expressed genes, we analyzed gene expression levels in genes hosting HIV-1 integration sites across five tissues: brain, colon, duodenum, esophagus, and stomach. Gene expression data were obtained from RNA-seq datasets available in the Human Protein Atlas. Genes were categorized into low-frequency integration (1–5 integration sites per gene) and high-frequency integration (≥6 integration sites per gene), and comparisons were made using a Mann-Whitney U Test (Wilcoxon Rank-Sum Test). Across tissues, the median gene expression levels showed variability between the low- and high-integration frequency groups, with some tissues exhibiting trends toward higher expression in genes with more integration sites (Figure 5D-H). However, statistical significance varied by tissue. In the colon, a significant difference in expression was observed (U = 521, p = 0.046), suggesting a potential preference for integration into highly expressed genes. In contrast, no significant differences were found in the brain (U = 333, p = 0.26), duodenum (U = 627, p = 0.10), esophagus (U = 949, p = 0.35), or stomach (U = 1249, p = 0.28).

Together, these findings reveal both tissue-specific and widely shared integration preferences, highlighting regions of the genome that may be particularly favorable for HIV-1 persistence. The presence of common integration hotspots across multiple tissues suggests that certain genomic loci could play a role in viral latency and reservoir maintenance.

### LTR Sequence Heterogeneity Confirms Biological Origins of Shared HIV Integration Sites

To assess whether PCR contamination contributed to the identification of shared HIV-1 integration sites across tissues and individuals, we analyzed the TBC1D5 hotspot, present in all tissues from all 13 PLWH. A total of 1,289 HIV-1 3’ LTR sequences integrated precisely at chr3:17443680 within the TBC1D5 gene were examined using multiple sequence alignment, mutation frequency analysis, and statistical comparison to known HIV-1 mutation rates. Additionally, we assessed host-mediated hypermutation signatures and searched for the TBC1D5 hotspot in independent datasets from the Retrovirus Integration Database (v2.0) (43).

Pairwise sequence alignments quantified nucleotide differences within the last 29 bp of the LTR, and a network-based analysis revealed high sequence diversity, with pairwise differences ranging from 1 to 5 bp (Figure 6A and 6B). While distinct sequence clusters were observed, no single dominant sequence indicative of widespread contamination was detected. Instead, several highly divergent sequences exhibited little to no connections to others, suggesting independent integration events rather than PCR amplification artifacts. If contamination had occurred, we would expect a single dominant sequence to appear across unrelated samples, along with reduced sequence heterogeneity. However, our results showed a naturally diverse sequence distribution, consistent with authentic HIV-1 integration rather than contamination. Furthermore, identical integration coordinates were associated with subtle but reproducible LTR sequence variations, a pattern expected from natural HIV-1 evolution rather than PCR artifacts. In addition, uninfected genomic DNA controls processed alongside the infected samples yielded zero integration sites, further ruling out contamination.

**Figure 6:**
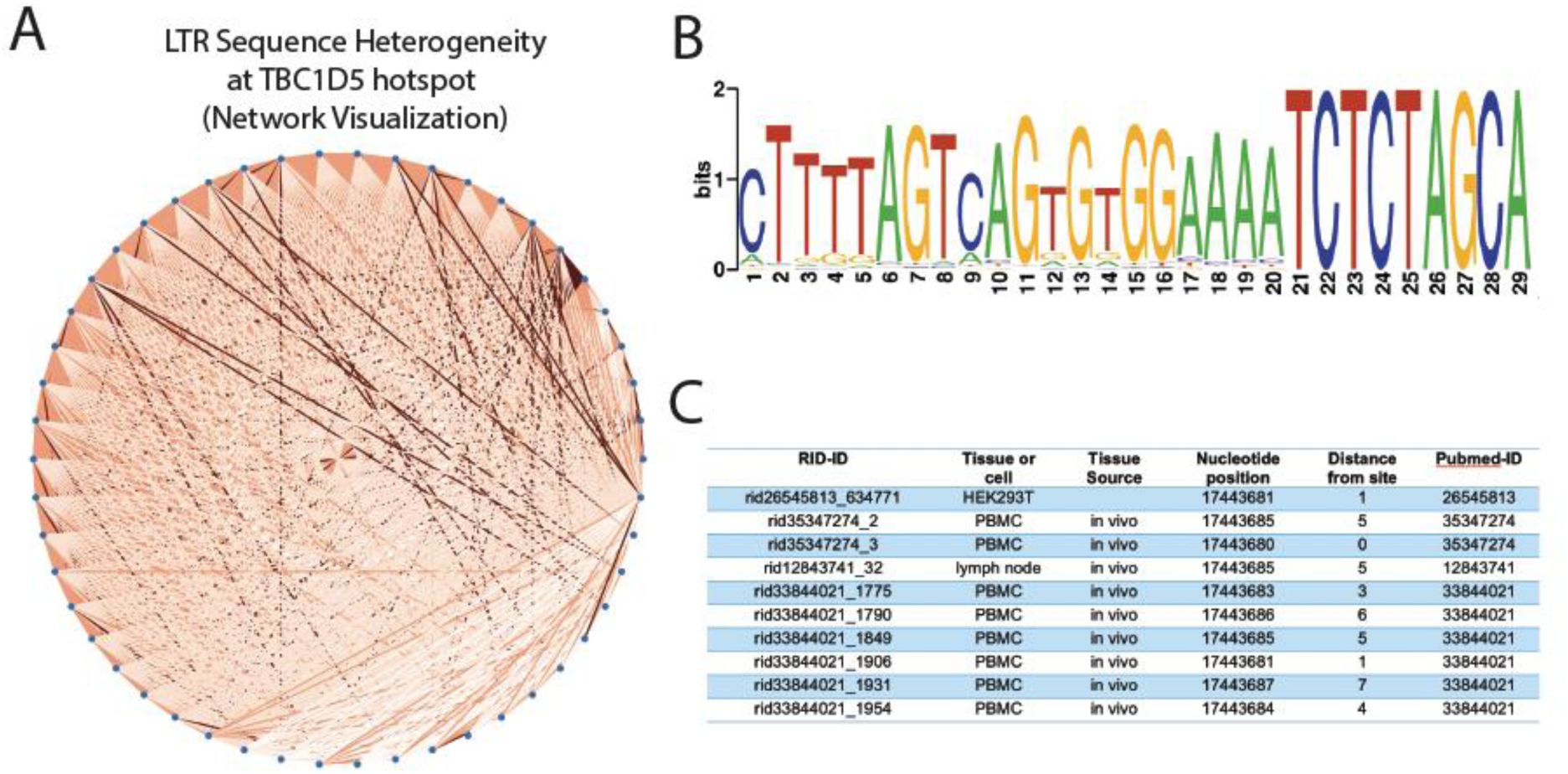
HIV-1 LTR heterogeneity at the TBC1D5 integration site hotspot. A, Network visualization representing the sequence heterogeneity among 1,289 HIV LTR sequences, focusing on the last 29 bp of the 3’ LTR region. Each node corresponds to an individual LTR sequence, while edges indicate pairwise nucleotide differences, with edge weight and color intensity reflecting the degree of divergence. Highly similar sequences form distinct clusters, while more divergent sequences exhibit fewer connections, suggesting independent integration events. B, Sequence logo depicting the nucleotide conservation and variability across the last 29 nucleotides of the HIV LTR region. The height of each nucleotide at a given position represents its relative frequency, with taller letters indicating higher conservation. C, Overlapping integration sites at the TBC1D5 integration site hotspot in independent integration site datasets available from the Retrovirus Integration Database (v2.0).

To determine whether the observed mutation rate aligned with known biological mechanisms, we compared it to expected HIV-1 RT and PCR polymerase error rates. Based on established RT mutation rates (2 × 10⁻⁵ to 5 × 10⁻⁵ per nucleotide per cycle), the expected mutation frequency for the 29 bp LTR region ranged from 0.00058 to 0.00145 per bp, with an expected G-to-A transition frequency between 0.000058 and 0.00022 per bp (44, 45). Similarly, the expected total PCR-induced mutation frequency was approximately 0.000725 per bp (high-fidelity polymerase, 1×10^−6^ errors per nucleotide for 25 cycles), with an expected 0.000073 per bp for G-to-A transitions (∼10%) (46, 47). A chi-square goodness-of-fit test revealed that the observed LTR mutation rate (0.034 per bp) was significantly higher than all expected background mutation rates (p < 0.0001), indicating that random RT and PCR errors alone cannot explain the observed sequence variability.

To evaluate APOBEC3-mediated mutagenesis, we quantified G-to-A transitions, a hallmark of APOBEC3 activity. These mutations accounted for 47% of all detected mutations (0.0162 per bp), a frequency significantly higher than expected RT (0.000058 to 0.00022 per bp) and PCR error rates (0.000073 per bp). A position-specific analysis identified 16 GG to AG mutations and 38 TC to TT mutations across 1,294 sequence variants. A chi-square test confirmed that these mutations occurred significantly more frequently than expected under a random mutation model (χ² = 8.96, df = 1, p = 0.00276). These findings provide evidence for APOBEC3-mediated G-to-A hypermutation, supporting the role of host antiviral defense mechanisms.

Finally, we examined whether the TBC1D5 hotspot was present in independent datasets from the Retrovirus Integration Database (v2.0). One PBMC dataset contained an integration site precisely at chr3:17443680, identical to our findings, and another integration site 5 bp away. Additionally, three other independent datasets involving HEK293T, PBMCs, and/or lymph node tissue showed integration sites 1 to 7 bp away from chr3:17443680, some of which were also shared between those independent datasets, reinforcing the biological relevance of this hotspot (Figure 6C). Together, our analysis demonstrates that the observed sequence diversity, mutation rates, and APOBEC3 mutation signatures cannot be attributed to PCR contamination. Instead, the results strongly support host-mediated mutagenesis and natural HIV-1 sequence evolution at the TBC1D5 integration site.

### HIV-1 integration targets genes implicated in diverse HIV-associated diseases

Proviral integration within or near genes can influence gene expression, potentially altering biological processes, molecular functions, and cellular pathways (reviewed in (48)). To investigate the relevance of integration site selection, we analyzed HIV-1 integration within genes linked to HIV-associated diseases using the Kyoto Encyclopedia of Genes and Genomes (KEGG) Disease database (Table 1). This analysis revealed striking tissue-specific integration patterns. Several genes implicated in cancer were frequent HIV-1 integration targets, with distinct patterns across tissues: brain, *PAN3*; esophagus, *CDK6*, *PRKG1*; stomach, *CBL*, *LDLR*; and colon: *BRIP1*, *KCNE2*. The gene *SCN5A*, linked to cardiovascular diseases, was a common integration target across all tissues. Additional cardiovascular-related genes were targeted in: esophagus, *PRKG1*; stomach, *LDLR*, *LMNA*; and colon, *KCNE2*. Several genes associated with nervous system diseases were highly targeted (>5 unique integration sites per gene): brain, *ATP6V0A1*, *ST3GAL3*; esophagus, *USP48*; stomach, *RIMS2*; duodenum, *DEPDC5*; and colon: *KCNT2*, *WNK1*. Several congenital malformation-associated genes were recurrent HIV-1 integration sites, including highly targeted genes in: brain, *ATP2C1*; stomach, *MACF1*; and duodenum, *MAPRE2*, *PPP2R5D*. HIV-1 integration was also observed in genes linked to a wide range of other diseases, including digestive, endocrine, hematologic, immune, metabolic, mental, respiratory, skin, urinary, and musculoskeletal disorders. This broad distribution suggests that integration site selection may contribute to the diverse health complications observed in HIV-infected individuals. Together, these findings highlight the potential impact of integration site selection on disease pathogenesis, emphasizing the need for further investigation into how proviral insertion influences gene function across different tissues.

### HIV-1 integration site hotspots across tissues are located in non-B DNA

Previous studies have shown that HIV-1 integration site hotspots are enriched in or adjacent to non-B DNA features, particularly slipped and G4 DNA motifs (28–30). To identify local genomic sequences that may serve as integration hotspots within tissues, we compared two pools of integration sites for each tissue: hotspot sites-integration sites occurring within hotspots (defined as two or more sites located <100 bp apart, regardless of genomic location); and non-hotspot sites-integration sites that did not cluster within hotspots. For each integration site, we extracted a 100 bp upstream and 100 bp downstream sequence window (200 bp total) and analyzed motif composition using DiffLogo, a tool that visualizes pairwise differences in DNA motifs (49). In DiffLogo visualizations, stack height represents the degree of sequence distribution dissimilarity (Jensen–Shannon divergence) and symbol height within each stack reflects the relative abundance of specific nucleotides at that position. Analysis of sequences surrounding integration sites revealed tissue-specific differences in motif composition (Figure 7). PBL/PBMCs exhibited the most sequence divergence, followed by colon, stomach, and duodenum. Esophagus showed the least sequence divergence, suggesting greater conservation of integration site sequences in this tissue. Consensus motif analysis across tissues revealed a consistent enrichment of slipped DNA motifs within hotspot regions (Figure 7). However, additional tissue-specific sequence preferences emerged. G4 DNA motifs were enriched at or near the integration site in esophagus, PBL/PBMCs, stomach, and duodenum. Brain tissue displayed overlapping STR and triplex motifs, positioned further downstream (66–87 bp from the integration site).

**Figure 7:**
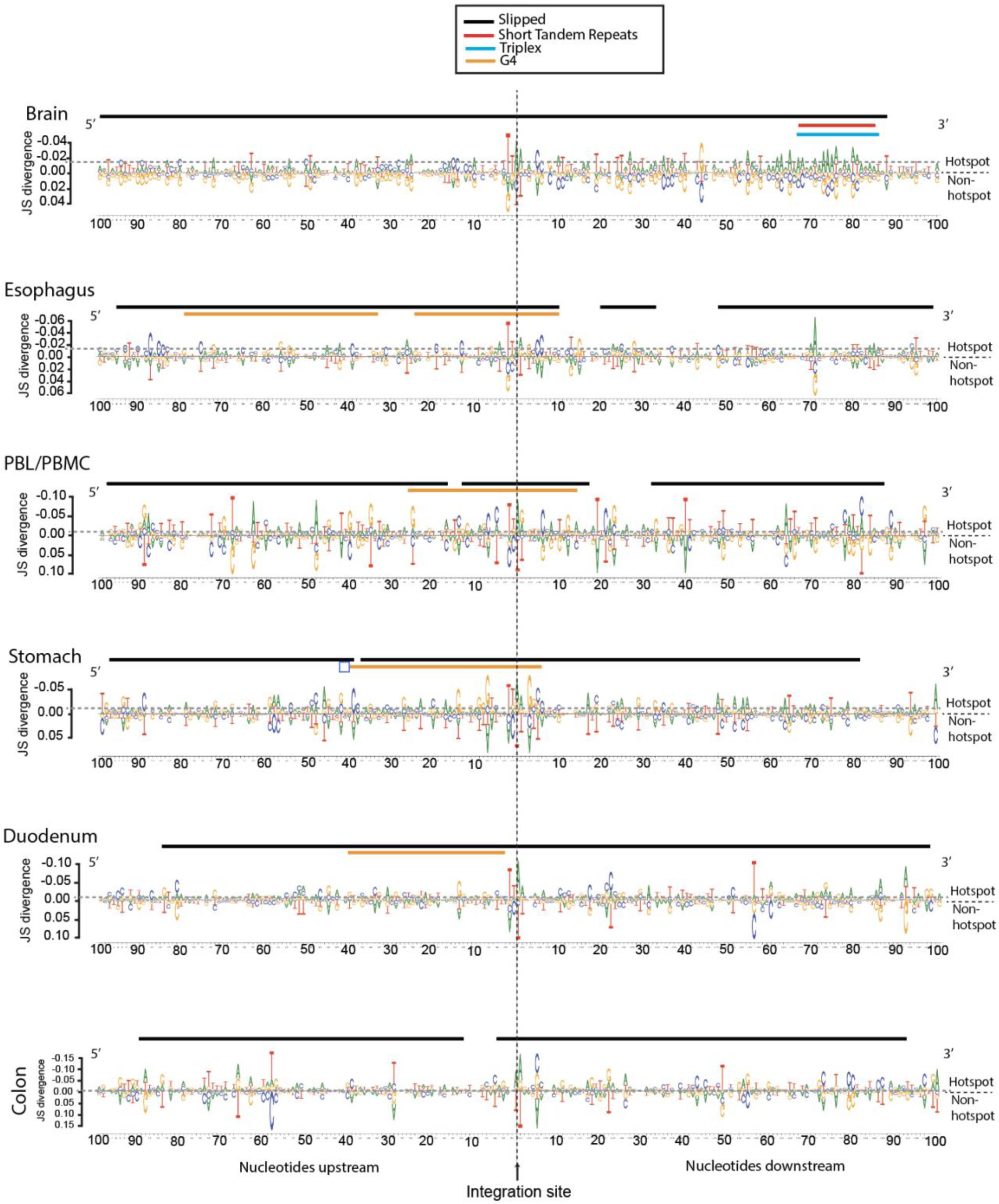
Integration site hotspots across tissues are located in non-B DNA. A, Genomic sequences were extracted from a window of 100 nucleotides upstream and 100 nucleotides downstream of each integration site. Sequences from integration sites located in hotspots were compared to sequences from sites not located in hotspots using DiffLogo. Consensus sequences were analyzed for the presence of non-B DNA motifs and represented by colored lines above each DiffLogo image (black, slipped DNA motif; orange, G4 DNA motif, red, STR; blue, triplex motif). The top half of each DiffLogo represents sequences from hotspots and the lower half represents sequences from non-hotspots. Vertical dashed line represents the point of integration; grey horizontal dashed lines represent JS divergence values of −0.02 as a reference point for comparison.

These findings underscore the conserved role of slipped DNA motifs in HIV-1 integration hotspots, while highlighting tissue-specific preferences for G4 motifs (esophagus, PBL/PBMCs, stomach, duodenum) and STR/triplex motifs (brain). Such sequence biases may influence regional chromatin accessibility and contribute to the establishment of tissue-specific viral reservoirs.

## DISCUSSION

This study provides new insights into HIV-1 integration preferences across anatomically diverse reservoirs. While most tissues (PBL/PBMC, esophagus, stomach, duodenum, colon) exhibited largely similar integration patterns with respect to common genomic features, brain tissue displayed a unique profile, with reduced gene targeting and increased integration into SINEs and DNase I hypersensitivity sites (DHS). Additionally, each tissue exhibited distinct preferences for integration near non-B DNA structures, and we identified integration hotspots across all tissues—some tissue-specific and others shared. Importantly, we observed substantial integration site overlap between tissues and individuals, suggesting highly precise targeting at specific genomic locations.

Understanding HIV-1 integration in the brain remains challenging due to limited tissue availability and the low frequency of infected cells in this compartment. A recent study in cultured microglia found that HIV-1 preferentially integrates into transcriptionally active chromatin and genes (50). Our finding that brain-derived integration sites were enriched in DHS aligns with this, as DHS mark regions of accessible chromatin, often associated with active transcription. However, brain tissue exhibited significantly less integration into genes than all other tissues, which was unexpected given the correlation between DHS and active transcription. A particularly striking finding was the increased integration into SINEs in brain tissue, while all other tissues showed reduced SINE targeting. SINEs (e.g., Alu repeats) and other transposable elements are commonly associated with gene silencing, as they are enriched in repressive chromatin marks such as histone H3 methylated at Lys9 (9, 10, 51). Although we did not differentiate between cell types in the brain, cell type-specific factors may influence integration preferences. One possible explanation is the role of Lens Epithelium-Derived Growth Factor (LEDGF/p75), a key host co-factor that directs HIV-1 integration toward active genes. LEDGF/p75 expression is regionally restricted and lower in the adult brain (52–58), potentially contributing to the reduced gene targeting observed. Conversely, APOBEC3G and APOBEC3F, which are known to shift HIV-1 integration away from genes and toward SINEs, are widely expressed in the central nervous system and can be upregulated by interferon (30, 59, 60). Together, decreased LEDGF/p75 and increased APOBEC3 expression may promote integration into transcriptionally repressed regions in brain cells.

A limitation of our study is that brain-derived integration sites were obtained from ART-naïve individuals, whereas integration sites from other tissues came from ART-treated individuals. This raises the possibility that the observed brain-specific integration patterns, such as reduced gene targeting and increased SINE enrichment, may reflect ongoing viral replication in untreated individuals rather than inherent tissue-specific factors. ART reshapes integration site landscapes by selectively expanding clonally integrated proviruses, particularly in actively transcribed genes (61). In untreated individuals, integration likely reflects initial infection dynamics, with broader distribution across genic and intergenic regions. This could explain the lower gene-targeting in brain tissue, as opposed to the gene-enriched profiles in ART-treated tissues, where clonal expansion dominates. Additionally, increased SINE integration in brain tissue may reflect a bias toward transcriptionally inactive regions, allowing proviruses to evade immune clearance. While our study does not directly compare pre- and post-ART integration landscapes, future research using matched pre-/post-ART samples or single-cell analyses of integration site clonality could clarify whether these differences are driven by intrinsic tissue properties or ART-related selection pressures. Recognizing this distinction highlights the need for further investigation into how ART shapes reservoir persistence across tissues, particularly in the brain.

Our analysis revealed substantial integration site overlap between different tissues within the same individuals. Given that all individuals in this study were MSM who likely acquired HIV-1 through direct gastrointestinal (GI) tract exposure, it is possible that infection was initially established in the lower bowel before disseminating to other GI sites and the bloodstream. This overlap could arise due to trafficking of infected cells containing identical proviruses, potentially due to clonal expansion, as observed in ART-treated individuals (40, 62, 63). Another possibility is migration of infected blood cells, leading to the seeding of identical proviruses across multiple tissues (31). Although our study did not assess clonality or proviral sequence identity, these mechanisms could explain the shared integration sites observed across tissues within PLWH.

Beyond intra-individual overlap, we also identified integration site overlap between different individuals, suggesting a common mechanism of integration site targeting during the early HIV-1 pandemic. Several factors could contribute to this: Transmission of a clonal HIV-1 strain—early in the pandemic, limited genetic diversity among circulating viruses may have led to similar integration patterns. A comprehensive sequencing analysis of the viral quasispecies from each patient would be needed to confirm this. Selective advantage—certain genomic regions may promote viral persistence or immune evasion, making them preferred integration sites across individuals. Host genetics—shared regions of DNA accessibility or chromatin structure may create similar integration site preferences. Convergent viral evolution—the selective pressure of early non-suppressive ART regimens may have driven the virus toward similar integration patterns over time. Further studies are needed to dissect how HIV-1 selects integration sites across different individuals and whether these patterns reflect evolutionary constraints, host factors, or a combination of both.

HIV-1 compartmentalization in CD4+ T cells across tissues suggests that integration site landscapes may differ between tissue-resident and circulating cells (31, 40, 41, 64). Some CD4+ T cell subsets remain tissue-resident or have limited ability to migrate, which could contribute to distinct integration site hotspots in different tissues. If ongoing viral replication occurs within these tissue compartments during ART, integration sites may continue to be generated at specific chromosomal regions that are highly receptive to integration. However, if there is no active viral replication during ART, as has been observed in lymphoid tissues (63), these integration events likely predate ART initiation and persist due to the long-lived nature of infected cells. Further investigation into proviral sequences, matching integration sites, and proviral RNA expression in tissues will help determine whether tissue-specific integration hotspots arise from ongoing replication or are remnants of past infection events.

The mechanisms that drive integration site hotspots, particularly those that are tissue-specific, remain poorly understood. Our previous work identified slipped DNA and G4 DNA motifs as common features of retroviral integration hotspots (28–30). In this study, we observed these same motifs within tissue-specific integration hotspots, suggesting that non-B DNA structures influence integration site targeting. Since slipped DNA and G4 DNA structures can be influenced by transcriptional activity, correlating host gene expression, proviral gene expression, and chromatin architecture with integration sites may provide key insights into why certain genomic regions are favored for integration in different tissues.

Retroviral integration near genes can disrupt gene function, leading to diverse biological consequences (48). While insertional mutagenesis has been implicated in the clonal expansion of infected cells, its role in HIV-1-associated diseases remains understudied. HIV-1-infected individuals face higher risks of AIDS-defining cancers (Kaposi sarcoma, aggressive B-cell non-Hodgkin lymphoma, cervical cancer) and non-AIDS-defining malignancies (65–68). While immune suppression and co-infections contribute to some of these cancers, the mechanisms underlying others remain unclear. Our discovery of HIV-1 integration within genes associated with various diseases, including cancer, cardiovascular disorders, and neurological conditions, raises an important question: Could HIV-1 insertional mutagenesis contribute to the wide range of clinical manifestations observed in people living with HIV-1? Determining whether HIV-1 integration into specific genetic loci drives disease progression will require further research, particularly longitudinal studies assessing gene expression changes in relation to integration events.

Despite providing valuable insights into HIV-1 integration site preferences across anatomically diverse tissues, this study has several limitations. First, the sample size is relatively small, particularly for brain tissue, which was obtained from an unmatched cohort. This limits the generalizability of our findings and may not fully capture the heterogeneity of HIV-1 integration patterns across different individuals and clinical backgrounds. Second, our analysis does not differentiate between specific cell types within each tissue, which is important given that HIV-1 exhibits cell type-specific integration preferences. For example, microglia and astrocytes in the brain may harbor HIV-1 differently than CD4+ T cells in the blood or gastrointestinal tract. Third, while we identified substantial integration site overlap across tissues and individuals, we did not assess whether these shared sites result from clonal expansion or other selective pressures. A deeper analysis of proviral sequence identity and transcriptional activity would help clarify the mechanisms underlying this overlap. Additionally, our study focuses on early-pandemic HIV-1 subtype B, and it remains unclear whether these tissue-specific integration site preferences extend to other HIV-1 subtypes or more current circulating HIV-1. Finally, while we identified associations between HIV-1 integration and specific genomic features, further mechanistic studies are needed to establish causal relationships between integration site selection, viral persistence, and disease outcomes. Future research incorporating single-cell sequencing, chromatin accessibility profiling, and host gene expression analyses will be critical for addressing these limitations and refining our understanding of HIV-1 reservoir dynamics across different tissues.

In conclusion, this study provides key insights into the complex relationship between HIV-1 integration preferences and tissue environments. We observed tissue-specific differences, with brain tissue exhibiting a unique integration profile. The identification of substantial integration site overlap and hotspots, including shared integration sites across individuals, suggests the presence of common targeting mechanisms. Further investigations across distinct cell types, including analysis of proviral sequences, transcriptional activity, and host epigenomic factors, will be essential to unravel the mechanisms driving tissue-specific HIV-1 integration and its role in disease development.

## Methods

### Sex as a biological variable

Patient samples were obtained from the cryobank without consideration of sex as a biological variable.

### Clinical Samples

Esophagus, PBL/PBMC, stomach, duodenum and colon tissue for this study was collected from five people living with HIV-1 (PLWH) as previously described (69–71). PLWH were enrolled from a cohort of HIV-1 seropositive men who have sex with men (MSM) followed at the Southern Alberta Clinic (SAC), Calgary, Alberta (Canada) between the years 1993 to 2010. Participants were prospectively followed and assessed for plasma viral load and CD4^+^ T counts were performed for each individual during each visit. Upper and lower gastrointestinal endoscopies were performed in order to collect biopsies of tissues from the esophagus, stomach, duodenum, and colon. Samples were cryopreserved during shipment and stored at −70°C within 1 hour of collection (69). PBL/PBMC were isolated from blood and stored in liquid nitrogen (71). The cohort from this study was recruited prior to the introduction of highly active antiretroviral therapy (HAART)/ART at the SAC. Participants received non-suppressive monotherapy or dual therapy with the nucleoside reverse transcriptase (NRTIs) such as azidothymidine (AZT/zidovudine), dideoxyinosine (ddI) prior to the study and during the study as previously described (71). Four participants also received HAART/ART at a later timepoint in treatment. Frozen brain samples originated from a separate unmatched cohort of HIV-1 seropositive patients not on ART and who succumbed to AIDS-defining illnesses (Table S1).

### Preparation of HIV-1 integration site library

Total genomic DNA was extracted from tissue samples using the DNeasy Blood & Tissue Kit (Qiagen). The purified genomic DNAs were processed for integration site analysis in a DNA clean room using different pipets, as previously described in detail (28–30). Briefly, the DNA was subjected to MseI/SacI di-gestion and linker ligation. Following purification, the DNA was subjected to two rounds of nested PCR using 3’ LTR- and linker-specific primers. The barcoded samples were sequenced through Illumina MiSeq using 2 × 150 bp chemistry at the London Regional Genomics Centre /Robarts Research Institute from Western University (Canada). Integration sites were determined from the sequence junction between the HIV-1 3’ LTR and human genome sequences.

### Computational Analysis

Each paired fastq sequencing read was quality trimmed and excluded from further analysis if the LTR-genome junction sequence did not match between the two paired reads. The HIV-1 LTR-containing fastq sequences were filtered by allowing up to a maximum of five mismatches with the reference LTR sequence. LTR sequences matching any region of the human genome (GRCh37/hg19) were discarded. Flanking human genomic sequences more than 20 nucleotides in length were used to identify integration sites using our in-house bioinformatics pipeline (Barr Lab Integration Site Identification Pipeline (BLISIP version 2.9)) (28–30). BLISIP version 2.9 includes the following updates: bedtools (v2.25.0), bioawk (awk version 20110810), bowtie2 (version 2.3.4.1), and restrSiteUtils (v1.2.9). All genomic sites within each dataset that hosted two or more sites (i.e., identical sites) were collapsed into one unique site for the analysis. Sites that could not be unambiguously mapped to a single region in the genome were excluded from analysis. All non-B DNA motifs were defined according to previously established criteria (72). Lamina-associated domains (LADs) were retrieved from http://dx.doi.org/10.1038/nature06947 (73). To account for restriction site bias in the cloning procedure during library construction, restriction enzyme site-matched random controls were independently generated for each dataset by matching each experimentally determined site with 50 random sites *in silico* that were constructed to be the same number of bases from the restriction site as was the experimental site, as previously described (74). The integration site heatmaps were generated using our in-house python program BLISIP Heatmap (BLISIPHA v1.0), which calculates the fold enrichment of sites in each distance bin for each feature compared to that of the matched random control dataset.

### LTR sequence heterogeneity

Raw HIV-1 LTR sequences were obtained from sequencing libraries prepared from multiple tissues and individuals. Quality control was performed to remove sequences with ambiguous bases (Ns) or deviations from the expected 29 bp length. PCR duplicates were filtered using USEARCH v11 (75). Sequences were aligned using MAFFT v7.490 in –auto mode with default gap penalties to preserve biologically relevant sequence variations (76). Pairwise nucleotide differences were calculated using a computational framework for sequence mutation analysis and visualized using NetworkX to generate a network-based representation of sequence relationships (77). Nodes represented individual LTR sequences, and edges indicated sequence differences, with edge weight and color intensity reflecting the degree of divergence. Only sequences with ≤5 nucleotide differences were connected to prevent network saturation. Mutation frequency analysis was conducted by calculating per-site mutation frequencies and comparing them to established HIV-1 RT mutation rates of 2 x 10⁻⁵ to 5 x 10⁻⁵ mutations per base per replication cycle (44, 45), or the error rate of high-fidelity DNA polymerases of 1 x 10⁻⁶ to 1 x 10⁻⁷ errors per base per replication cycle (46, 47). The expected mutation frequency per 29 bp LTR was derived and compared to observed mutation rates using a chi-square goodness-of-fit test with a significance threshold of α=0.05 was used. Additionally, the presence of APOBEC3-mediated hypermutations was evaluated using a computational framework for sequence mutation analysis. G to A transition frequencies and specific APOBEC3 target motif mutations (GG to AG and TC to TT transitions) were identified. The statistical enrichment of these motifs was assessed by comparing observed vs. expected mutation frequencies, using background HIV-1 mutation rates from published studies (78).

### Study Approval

Ethical approval for use of human esophagus, PBL/PBMC, stomach, duodenum and colon tissue were obtained from the Conjoint Health Ethics Research Board (CHREB, protocol approval #: REB15-1941) at the University of Calgary (Calgary, Alberta, Canada). Ethical approval for use of human brain tissue was obtained from the University of Alberta Human Ethics Committee, protocol approval #: Pro00002291. Participants signed an informed consent upon enrollment.

## Data availability

Data is available in the supplementary data files. Additional data is available upon request.

## Author Contributions

Research design: H.K., H.A., S.B., G.vM; Conducting experiments: H.K., H.A., E.A.B., N.R.V., I.K.F.W, M.C., S.T., S.B.; Acquiring data: H.K., H.A., E.A.B., N.R.V., I.K.F.W, M.C., S.T., S.B.; Analyzing data: H.K., H.A., J.G., D.C., P.B., C.P., G.vM., S.B.; Providing reagents: S.B., G.vM.; F.vdM., J.G., D.C., P.B., C.P.; Writing the manuscript: H.K., S.B.; Editing the manuscript: S.B., G.vM.; J.G., D.C., P.B., C.P.

## Acknowledgments

Funding was obtained from: Canadian Institutes of Health Research (CIHR) (FRN-150406) to S.B.; National Health Research and Development Program (NHRDP) (Grant No. 6609-1782-AIDS) to D.C.; Canadian Institutes of Health Research (CIHR) (CIHR-III HIV/AIDS HOP-9018) to G.vM.

